# Synthesizing Combination Therapies for Evolutionary Dynamics of Disease for Nonlinear Pharmacodynamics

**DOI:** 10.1101/009480

**Authors:** Vanessa Jonsson, Nikolai Matni, Richard M. Murray

## Abstract

Our previous results proposed an iterative scalable algorithm for the systematic design of sparse, small gain feedback strategies that stabilize the evolutionary dynamics of a generic disease model with linear pharmacodynamics. In this manuscript, we use piecewise linear approximations to model nonlinear drug effects. We leverage results from optimal controller synthesis for positive systems to formulate the feedback synthesis problem as an optimization problem that sequentially explores piecewise linear subsystems corresponding to higher and higher treatment dosages.

## I. INTRODUCTION AND MOTIVATION

A challenge inherent to the treatment of certain infectious and non-infectious diseases, such as HIV or cancer, is the risk that the pathogen or tumor will evolve away and become resistant to treatment methods that comprise the standard of care [1], [2], [3], [4]. Especially vulnerable to this phenomenon are treatment methods that involve exposing the disease population (such as viruses or cancer cells) to therapies targeting specific molecules involved in disease progression for an extended period of time. While these targeted therapies have the benefit of allowing physicians to tailor treatments to a patient’s tumor cell population, they nonetheless establish an environment in which the occurrence of mildly drug resistant pathogens or tumor cells can develop an evolutionary advantage over those for which the therapy is targeted [5], [6], [7], [8], leading to so called ‘treatment-escape’.

Pharmacodynamic models are used to quantitatively describe nonlinear drug dose-responses and model drug-receptor relationships. These models are often described using combinations of nonlinear Hill functions, and allow for the modeling of drug saturation effects [9], [10], [11]. These pharmacodynamic nonlinearities are further increased with the fact that drugs administered in combination can have independent additive effects or can otherwise exhibit synergistic or antagonistic dose dependent behavior, further complicating the design of suitable combination therapies [12].

The challenge of designing treatment protocols that prevent escape is one that has been addressed by control theoretic methods. For cancer therapy, results in this spirit apply methods from optimal and receding horizon control [13], [14], as well as gain scheduling [15], to synthesize treatment protocols that are robust to parameter uncertainty, an inherent issue in all biological systems. The authors in [16], present a static multi-objective optimization formulation to solve the combination therapy problem for different initial tumor populations, when the drugs under consideration have additive, linear effects on cell viability. Proposed combination treatments were confirmed experimentally for different tumor initial conditions in a murine lymphoma model [17]. In the context of HIV and antiretroviral therapy, the authors in [18] propose a discrete time formulation that allows for the design of switching therapy strategies to delay the emergence of highly resistant mutant viruses. Recent results in [19] and [20], consider a simplified bilinear model and the optimal control problem is shown to be convex over a finite horizon for a predefined set of initial states.

There have been several attempts to deal with nonlinear HIV infection dynamics using model predictive control (MPC) to design optimal antiretroviral drug dosing strategies [21], [22]. An issue with the use of MPC is that an essential feature of the system, that of nonlinearities associated with drug binding, is linearized away. In particular, such an approach can lead to a model that underestimates the efficacy of a drug at lower concentrations, and over estimates its efficacy at high dosages *unless* a sufficiently small update time is taken. This latter restriction may then lead to strategies that are no longer realistic in a biomedical application, where it may be difficult to update the administered therapies at the frequency dictated by the MPC controller. In this paper, we take an alternative approach and attempt to design constant drug therapies by taking input non-linearities into account more explicitly.

In [23], we introduced a scalable, iterative algorithm for the principled design of targeted combination therapy concentrations that explicitly accounts for the evolutionary dynamics of a generic disease model. This algorithm was effective in generating robustly stabilizing controllers, while simultaneously allowing the designer to explore explicit trade offs between closed loop performance, sparsity in controller structure and gain minimization. Leveraging recent results from positive systems [24], we formulated this algorithm as a second order cone program (SOCP), which made the controller synthesis scalable. However it could not take into account a) the pharmacodynamics of the input, potentially suffering from over or underestimating gains, and b) the effects of synergistic or antagonistic drug combinations that can be modeled with additional nonlinear pharmacodynamic terms [12].

Here we propose a new algorithm that solves the combination therapy problem subject to the same design constraints (sparsity of the drug combination, maximum dosage and robustness constraints) formulated as an SOCP while addressing the non linear dynamics of individual drugs and of their combinations. In particular, through the piecewise linearization of individual and combination drug pharmacodynamics, in conjunction with a branch and bound like algorithm for the effective search through these linear pharmacodynamic modes, we reduce the combination therapy problem to applying the SOCP formulation from [23] to a set of pharmacodynamic modes.

The main contribution of this paper is an algorithm for the systematic design of *sparse*, *small gain* feedback strategies to stabilize the evolutionary dynamics of a generic disease model and general nonlinear pharmacodynamics models, which support synthesis of feedback strategies in light of highly nonlinear drug dynamics.

The article is structured as follows: In Section II, we recall the extended quasispecies evolutionary dynamics model that encodes replication, mutation and neutralization and summarize relevant results in controller design of positive systems and introduce the pharmacodynamics of a non competitive additive drug binding model. In Section III, we present our 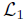 combination therapy synthesis algorithm as well as an algorithm designed to increase the scalability of the formulation. Section IV illustrates our algorithm in the context of an additive drug interaction example, that of the HIV antibody therapy design problem previously studied in an experimental setting [25], where the drugs act independently and additively. Section V ends with concluding remarks and directions for future work.

## II. PRELIMINARIES

### A. Notation

ℝ_+_ denotes the set of nonnegative real numbers. The inequality *X* > 0, (*X* ≥ 0) means that all elements of the matrix or vector X are positive (nonnegative). *X* ≻ 0 means that *X* is a symmetric and positive definite matrix. The matrix *A* ∈ ℝ*^n^*^×*n*^ is said to be Hurwitz if all eigenvalues have negative real part. Finally, the matrix is said to be Metzler if all off-diagonal elements are nonnegative. Define 1*_n_* to be the vector of all ones of dimension *n*. The induced matrix norm for a matrix *M* ∈ ℝ^*r*×*m*^ is 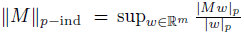 where 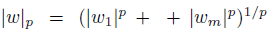. Let *G*(*s*) = *C*(*sI* – *A*)^−1^ *B* + *D* be a *r* × *m* matrix transfer function. The induced norms of the corresponding impulse response *g*(*t*) = *Ce ^At^B* + *Dδ*(*t*) are 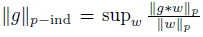 for 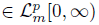, given that 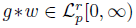 is the convolution of *g* and *w*. Finally we refer to the ∞-induced robust controller as the 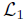 controller as is customary in the robust control literature.

### B. Evolutionary dynamics model

The quasispecies model [26] was originally developed to describe the dynamics of populations of self replicating macromolecules undergoing mutation and selection. We choose this model for its relative simplicity and its ability to capture the salient features of the evolutionary dynamics of a simplified generic disease model. In [27] we incorporated the effects of potential therapies into the basic quasi species model, by defining a drug binding reaction, 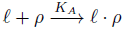 — drug *ℓ* binds to self replicating macromolecule *ρ* with association rate *K_A_*, giving a neutralized complex *ℓ* · *ρ*. The extended quasispecies model for *n* mutants and *m* drugs, is written as:

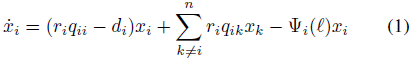

where *x_i_* ∈ ℝ_+_ is the concentration of mutant *i*, 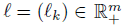 is the drug concentration (assumed to remain at constant concentrations throughout), *r_i_* and *d_i_* are the replication and degradation rates, respectively, of mutant *i*, and *q_ik_* is the probability that mutant *k* mutates to mutant *i* (note that *q_ii_* is the probability of no mutation occurring). The rates *r_i_* can be viewed as the replication fitnesses of mutant *i* without the effect of the drug. When *ℓ_k_* = 0, ∀*k* ∈ {1*,…, m*}, the quasispecies dynamics are unstable. Finally, the function Ψ*_i_* (*ℓ*) represents the pharmacodynamics of individual drugs *ℓ_k_* and their combinations with respect to the *i*-th mutant species, namely the sum of nonlinear drug effect functions as represented by Hill equations. In the following section, we present a brief justification for the Hill equation models of these binding reactions.

### C. Pharmacodynamic models

When combined, drugs can have additive, synergistic or antagonistic effects that need to be explicitly accounted for when designing combination therapies. When the presence of one drug modulates the effect of another, the combined drug effects are no longer additive, and in particular, the phamacodynamics of the drug interaction now incorporate additional terms that represent this interaction.

#### 1) The Hill equation

The Hill equation has been used in pharmacology to model nonlinear drug dose-responses, for example the effects of drugs on cell viability or virus neutralization. More generally it serves to quantify drug-receptor relationships, the fraction of bound receptors *ρ* (e.g. cell receptors, virus proteins) as a function of ligand *ℓ_k_* (e.g. drug, antibody) concentrations for the binding reaction

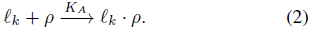

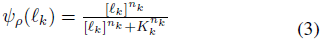

where *ψ_ρ_*(*ℓ_k_*) ∈ [0, 1] is the fraction of bound receptors, *ℓ_k_* ∈ ℝ_+_ is the concentration of ligands, 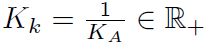 is the dissociation constant associated with the binding reaction (4), *n_k_* ∈ ℝ_+_ is the Hill coefficient that represents the degree of cooperativity, i.e. the degree to which binding of a ligand molecule modulates the probability of another ligand molecule binding.

We notice that the Hill function is a biological analog to actuator saturation, in that there is a law of diminishing returns in terms of the effect of ever increasing drug concentrations on the system. In fact, the Hill functions *ψ_ρ_*(*ℓ_k_*) that appear in HIV applications, for example, look approximately linear for small to moderate *ℓ_k_*, and nearly constant for large *ℓ_k_* – it is this observation that motivates our piecewise linear approximation approach.

#### 2) Non-competitive additive drug binding model

Under this model, we consider the pharmacodynamics of non interacting drugs that bind to different receptors, in which the effect of each drug on the system is additive. Consider the system of *m* drug binding reactions to different receptors on a particular cell or virus *x*:

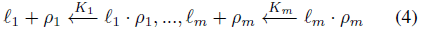

where *ℓ_k_* is the drug *k* and *ρ_k_* is a receptor. If these receptors comprise different drug binding targets on a cell or virus *x*, then we can describe the total effect of these independently acting drugs *ℓ*_1_, …, *ℓ_m_* on *x* as

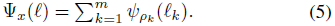

##### Remark 1

Although the remainder of the paper focusses on this drug interaction model, we note that our approach applies nearly verbatim to a synergistic/antagonistic drug binding model – it suffices to use a suitable expression for Ψ*_x_*(*ℓ*) to take these interactions into account. We do note however that this may lead to a more involved piecewise linear approximation procedure.

## III. A SUB-OPTIMAL 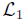 COMBINATION THERAPY CONTROLLER FOR NON LINEAR PHARMACODYNAMICS

In this section, we formulate the task of designing suitable combination therapies as an optimal control problem. The inherent non-linearities of the system make this a challenging task – in [23], [27], we worked with a simplified problem in which we assumed the Ψ*_x_*(*ℓ*) were linear functions – in this paper we relax that assumption, and show that at the expense of some additional modeling complexity, we are able to reduce the problem to that considered in [23]. Our main result is based around the use of piecewise linear approximations, and relating the robustness levels of the approximate system to that of the true underlying system.

### A. Problem formulation

The following state space representation of equation (1) emphasizes the inherent feedback structure that arises from drug binding reactions:

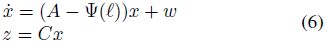

with (i) *A* ∈ ℝ^*n×n*^, with *A_ij_* = *r_i_q_ij_* ≥ 0 ∀ *i* ≠ *j* and *A_ii_* = *r_i_q_ii_* – *d_i_*, that encodes the mutation and replication dynamics; (ii) Ψ(*ℓ*) ∈ ℝ^*n×n*^, a diagonal matrix, with diagonal elements Ψ*_xi_* (*ℓ*), that maps the effects of the therapies *ℓ* to the population dynamics; (iii) *ℓ* = (*ℓ_k_*) ∈ ℝ^*m*^, a vector of the concentrations of neutralizing macromolecules; (iv) 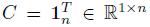; and 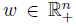 an arbitrary positive disturbance.

We set the regulated output *z* = 1*_n_x* to be the total virus population, so as to ensure that the resulting treatment plan is one that robustly drives the total mutant population to zero.

#### Remark 2

*A* is a Metzler matrix with off-diagonal entries that are several orders of magnitude smaller than the diagonal entries. This is due to the biological fact that mutation rates range from 10*^−^*^5^–10*^−^*^8^ mutations per base pair per replication cycle for reverse transcriptase [28] to DNA replication [29].

Letting *G* denote the closed loop system (6), the control task then becomes to reverse engineer neutralizing macro-molecule concentrations *ℓ* by finding a ‘‘controller” Ψ(*ℓ*) that leads to a stable *G* satisfying 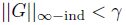, for some robustness level γ > 0.

### B. Piecewise linear mode approximations and mode reduction

In the following, we assume that each (Ψ(*ℓ*))*_ii_* = Ψ*_xi_* (*ℓ*) has the form given by (5). To take into account non-linear pharmacodynamics, we propose a piecewise linear approximation algorithm and a mode reduction algorithm for problems where there is a large number of non-interacting or synergistic drug combinations.

We assume that the pharmacodynamics for every individual drug and combination are defined over the same drug concentration domain 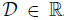. Let 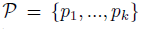 be a partitioning of this domain into *k* intervals.

#### Definition 1

A pharmacodynamic mode *ω* = (*ω*_1_*,…, ω_m_*) is an *m*-tuple in 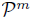. The total number of pharmacodynamic modes is 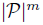 where *m* is the number of drugs under consideration. For *ℓ* ∈ ℝ^*m*^, we define 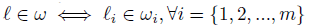.

We let *ψ_iω_*: ℝ^*m*^ → ℝ be the affine approximation of the pharmacodynamics for each mutant *i* for *ℓ* ∈ *ω*, i.e. the sum of the individual and combination drug effects on mutant *i* while operating within mode *ω*. We can then construct a linear approximation to Ψ(*ℓ*) via an appropriately defined block diagonal matrix Ψ*_ω_*, constructed from the *ψ_iω_* (c.f. [23]), and write Ψ(*ℓ*) = Ψ*_ω_L_ω_*, where 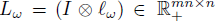 is the block diagonal matrix, with identical block diagonal elements given by the drug concentrations *ℓ* ∈ *ω*. The resulting dynamics, for a fixed concentration *ℓ* ∈ *ω*, are then described by the transfer function

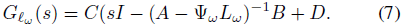

We consider the problem of finding a suitably sparse therapy combination that achieves a certain closed loop performance level *γ*. As such, our initial goal becomes to reduce the search space to the set of sparse modes that achieve the desired level of robustness, where a sparse mode *ω* is one that allows at least one drug concentration to be zero, in other words, 0 ∈ *ω_i_* for at least one *ω_i_* ∈ *ω*.

In order to do so, we require two lemmas. The first provides sufficient conditions on the linear approximation terms *ψ_iw_* that guarantee that the robustness of the piecewise linear approximation is an upper bound on that of the true system. The second is the simple observation that for non-interacting or synergistic additive drug interactions, the robustness of the closed loop dynamics increases as drug concentrations are uniformly increased (this statement will be made precise). This result allows us to subsequently develop a branch and bound like method that significantly reduces the search space of the algorithm.

We begin with a result on the input-output performance of a positive system, taken from [24].

#### Lemma 1

Let *G*(*s*) = *C*(*sI − A*)*^−^*^1^*B* + *D* be a positive system. Then 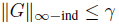 if and only if there exits *x ≥* 0 such that

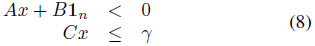

#### Lemma 2

Let Ψ(*ℓ*) be the non-linear pharmacodynamics function, *ψ*(*ℓ*) the vector of its diagonal elements, and denote by Ψ*_ω_*(*ℓ*) its piecewise linear approximation within mode *ω*, and by *ψ_ω_*(*ℓ*) the vector of its diagonal elements. If for every mode *ω* we have *ψ_ω_*(*ℓ*) ≤ *ψ*(*ℓ*) ∀*ℓ* ∈ *ω* then the 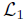 norm *γ* of the piecewise linear approximation (7) is an upper bound on that of the true system (6).

*Proof:* Note that for a fixed *ℓ*, both the full and piecewise linear approximation systems are linear in *x*. Therefore, by Lemma 1, the 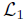 norm of (7) is upper bounded by *γ* if and only if there exists an *x >* 0 such that

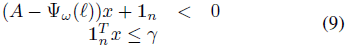

Thus it suffices to show that this same *x* yields (*A −* Ψ(*ℓ*))*x* + 1*_n_* ≤ 0 and the desired conclusion follows immediately. To that end, rewrite the first inequality of (9) as

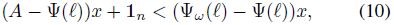

and notice that the right hand side is less than or equal to 0 by the assumptions of the lemma.

This lemma essentially states that if our piecewise linear approximation is conservative, then the norm of the approximate system serves as a certificate for the norm of the true system.

Next we formalize the observation that increasing the concentrations of non or synergistically interacting therapies present in the system will improve robustness.

#### Lemma 3

Let *ℓ*^1^ and *ℓ*^2^ be therapy combinations such that *ℓ*^1^ *≥ ℓ*^2^. Then if the piecewise linear approximation Ψ*_ω_*(*ℓ*) is non-decreasing, 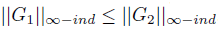, where

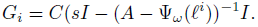

*Proof:* By assumption, the piecewise linear function Ψ*_ω_*(*ℓ*) is non-decreasing. Thus if *ℓ*^1^ *≥ ℓ*^2^, then Ψ*_ω_*(*ℓ*^1^) *≥* Ψ*_ω_*(*ℓ*^2^). Let 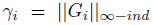. By Lemma 1, *γ_i_* is the solution to the following optimization:

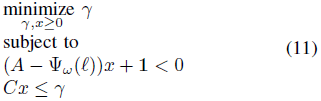

Let *γ*_2_ and *x* be the optimal solutions of the above program for *i* = 2. Then we have that

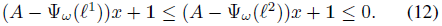

Hence (*γ*_2_*, x*) is a feasible solution for optimization (11) with *i* = 1, implying that *γ*_1_ *≤ γ*_2_.

Thus we see that by constraining the piecewise linear approximations to be under approximations of the effects of the drugs, we are able to bound the performance of the true system.

#### Remark 3

We note that these results typically will not hold for antagonistic drug interactions – however, many settings in which such models are required (such as cancer therapy design) do not have a large set of therapies or mutants, thus mitigating the computational cost of the sequential search across modes.

We exploit this result to reduce the modes that need to be searched over – in particular, we use the partial order implied by the previous two results to upper and lower bound the uniform concentration treatments required to achieve a prescribed performance level *γ*. We also exploit the fact that we are searching for *sparse* treatment strategies to further eliminate modes.

This approach is formalized in the following algorithm. Let *ℓ_ω_max__* ∈ ℝ*^m^*, be the maximum possible drug concentrations.

#### Algorithm 1 Sparse mode reduction algorithm

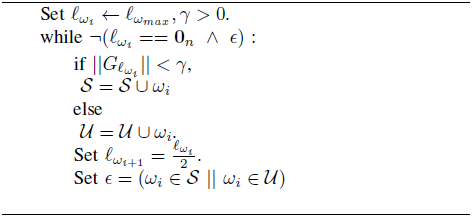

The sparse mode reduction algorithm will generate a set of modes that are guaranteed to be stable and achieve a desired robustness level *γ*, and “sparse”, i.e. allowing modes such that at least one drug concentration is allowed to be zero, significantly reducing the number of the modes over which to apply the combination therapy algorithm. For the HIV example in Section IV, to synthesize controllers with robustness level *γ* = 14, we start with 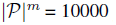 possible modes to search over, and reduce this number to 397 sparse modes with robustness level *γ* = 14, via Algorithm 1.

#### Algorithm 2 Scalable Combination Therapy

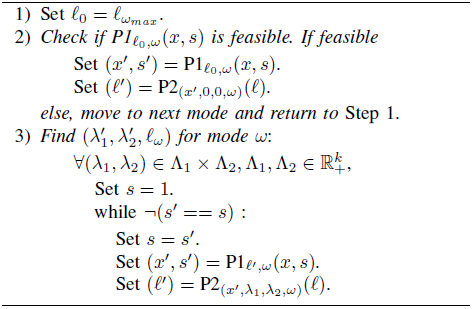

### C. A suboptimal combination therapy algorithm

As discussed in [23], [27] there are no known convex reformulations for the robust combination therapy problem over an infinite horizon due to the additional structure on *L*. As such we use the iterative approach developed in [23], as formalized in Algorithm 2, based on the convex programs (13) and (14), to find a stabilizing controller, given a desired robustness level *γ*. For notation, let *Y*′ = P*_Z_*_′_(*x, s*) denote an optimization problem P in which we optimize over *x* and *s* leaving *Z*′ fixed and with solution *Y*′. These optimization programs, taken from [24], are a synthesis variant of the conditions stated in Lemma 1.

#### Program 1.

P1_*ℓ*_*_,ω_*(*x, s*) :

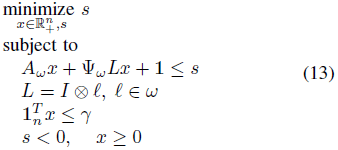

#### Program 2.

P2_(*x*,λ1, λ2, ω)_(*ℓ*)

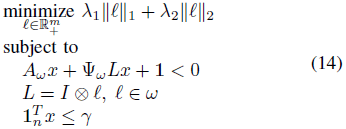

#### Remark 4

Note the introduction of a slack variable *s* into Program 1, to help prevent immediate convergence to a local minimum. Minimizing *s* has the effect of maximizing the slack in the first constraint, allowing for more freedom in the design of the concentration vector *ℓ* in Program 2.

## IV. APPLICATIONS TO ADDITIVE PHARMACODYNAMIC BINDING MODELS

### A. Additive drug effects: HIV and antibody therapy

Our results provide a principled approach to the design of antibody treatments for chronic infection with human immunodeficiency virus-1 (HIV-1) in light of the nonlinear pharmacodynamics and saturation concerns associated with antibody neutralization of HIV-1. We illustrate this with an example motivated by experimental results of evolutionary dynamics of HIV-1 in the presence of antibody therapy obtained in [25].

A relatively recent discovery is that a minority of HIV-infected individuals can produce broadly neutralizing antibodies (bNAbs), that is, antibodies that inhibit infection by many strains of HIV [30]. These have been shown to inhibit infection by a broad range of viral isolates *in vitro* but also protect non-human primates against infection [30],[31], [32]. Recent experimental results conducted in the Nussenzweig lab at Rockefeller University have demonstrated that the use of single antibody treatments can exert selective pressure on the virus, but escape mutants due to a single point mutation can emerge within a short period of time [25]. Although antibody monotherapy did not prove effective, it was shown that equal, high concentrations of an antibody pentamix effectively control HIV infection and suppress viral load to levels below detection.

The goal of this example is to demonstrate how our proposed algorithm offers a principled way to design combination antibody therapies that control HIV infection and prevent evolution of any set of known resistant mutants given the nonlinear pharmacodynamics of antibody neutralization of HIV-1. In a realistic setting, the ability to do this relies on the knowledge of what resistant viruses may be selected for with single therapies, and knowledge of antibody pharmacodynamics. This algorithm would be most effective in conjunction with single antibody selection experiments and knowledge of antibody Hill function properties.

#### 1) Model parameters

We consider four potential antibodies to use in combination PG16, 45-46G54W, PGT128 and 10-1074 on an evolutionary dynamics model of twenty one mutants.

The Hill function associated with the fitness of a virus with respect to neutralization by an antibody is described in equation (3). We used Hill function parameters that were experimentally derived in [33] for antibodies 45-46G54W, PGT128 and PG16 and approximated the Hill parameters associated with antibody 10-1074. We consider a set of twenty one mutants that evolved from antibody monotherapy experiments in [25]. Their corresponding half maximal inhibitory antibody concentration (IC50) in *µg/ml*, is specified in the Supplementary Figure 8 of [25]. We choose replication rates to be 0.5 (ml · day)*^−^*^1^ for all mutants. We justify this selection by noting that escape mutants grew to be dominant mutants during selection experiments and assume that replication rate variability due to mutations are negligible. The mutation rate for HIV reverse transcriptase is *u* = 3 × 10*^-^*^5^ mutations/nucleotide base pair/replication cycle [28], and the HIV replication cycle is approximately 2.6 days [34]. We approximate the rate of mutation for a particular amino acid mutation at a particular location to be 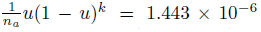 per replication cycle, where *k ≈* 3000 is the size of the genome in residues and *n_a_* = 19 is the number of amino acids that can be mutated to. Our model supports forward point mutations and two point mutations. We do not consider back mutations, as the probability of mutation is negligible. Units of concentration in number of viruses/ml or number of antibodies/ml are used for states, and time is measured in days. The standard volume is 1 ml.

### B. L_1_ controller synthesis

We performed ten piecewise linear approximations on each of the Hill functions associated to each of the twenty one mutants and each of the four antibodies we considered as possible candidate therapies. This generated 10000 possible pharmacodynamic modes of which we found 397 to be sparse and stable by Algorithm 1. Figure 1 shows an example of a conservative piecewise linear approximation to a Hill function associated with a virus mutant antibody pairs.

**Fig. 1.**
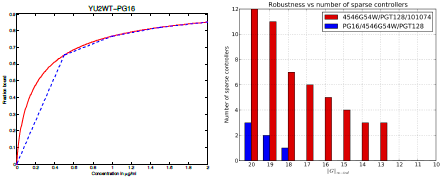
(Left) Piecewise linear approximation (blue dashed) of the Hill function (red) for PG16-wild type clade B YU2 HIV-1. (Right) Robustness versus number of sparse controllers synthesized using the 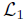 nonlinear combination therapy algorithm for the first lexicographically ordered twenty robustly stable sparse modes. Two different 3-sparse antibody trimixes are synthesized, with the dominant trimix being the one that excludes PG16.

We synthesized a family of robustly stabilizing controllers using our algorithms for a range of desired robustness levels and found that two different trimixes were predominant - one more frequent combination comprised of (45-46G54W, PGT128,10-1074) antibodies was present for smaller robustness levels and another combination, (PG16, 45-46G54W, PGT128) appeared in addition to the first one when the desired robustness was allowed to be larger as shown in Figure 1. According to [25], the mean IC50 for virus mutants is greatest for the PG16 mono therapy than any other single antibody.

In [25], a different antibody trimix containing PG16, (3BC176, PG16, 45-46G54W) was suggested and experimentally shown to produce a decline in the initial viral load. However, a majority of mice in the experimental study had a viral rebound to the trimix pre-treatment levels, suggesting that in these cases, the virus had evolved mutations that were resistant to the trimix treatment. Further study showed that the evolved mutants had mutations found in the PG16 and 45-46G54W monotherapy groups. This could be suggestive that the combination of (PG16 and 45-46G54W) with another antibody although stabilizing, may not be robust enough to the type of perturbations witnessed in a biological setting. In Figure 2, we see that although the (PG16,45-46G54W, PGT128) (red) and (45-46G54W, PGT128,10-1074) (green) and combinations were synthesized to have the same robustness guarantee, the combination containing PG16 and 45-46G54W is less robust to perturbations.

**Fig. 2.**
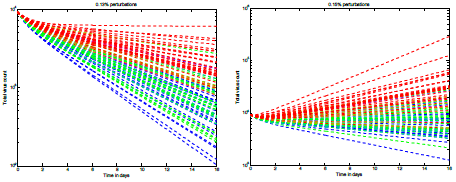
Sum of virus populations subject to random time invariant perturbations of 0.13% in the dynamics for 30 different simulations for (left) a stabilizing closed loop controller comprised of antibody trimixes (0,0.96,2,1) (blue), (0,1,0.177,2) (green) and (2,0.35,0.96,0) (red) *µg/ml* of (PG16,4546-G54W, PGT128, 10-1074) combinations synthesized using our algorithm and (right) for the same antibody trimixes subject to perturbations of 0.15% in the dynamics.

These simulations demonstrate that although many stabilizing solutions to the combination therapy problem exist, the best ones are found when design parameters such as a sparsity, limits on the magnitude of gains, and robustness guarantees are simultaneously considered. Experimentally searching for these combinations is infeasible as the number of potential therapies and possible concentrations to consider is experimentally intractable. We propose to guide these experimental activities with our ability to design and synthesize combination therapy controllers. As such, one could generate a family of controllers based on “design specifications” tailored not only the (viral or cellular) composition of the disease, but to explore tradeoffs between number of therapies used (sparsity), therapy concentrations (magnitude of the gain) and ability to support pharmacokinetic fluctuations (robustness to perturbations) and subsequently verify these experimentally.

## V. CONCLUSION AND FUTURE WORK

We proposed an iterative algorithm and mode reduction scheme for the systematic design of sparse, small gain feedback strategies that stabilize the evolutionary dynamics of a generic disease model, subject to nonlinear pharmacodynamics. In future work, we plan to explore a principled integration of our methods with recent results on the robust 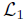 stability of positive systems [35]. In particular, we hope to explicitly account for unmodeled dynamics due to pharmacokinetic perturbations.

## VI. ACKNOWLEDGEMENTS

We would like to thank David Baltimore and Pamela Bjorkman for discussions regarding using antibody therapy for HIV treatment. We appreciate the help of Bjorkman laboratory staff scientist Anthony West for information on antibody neutralization parameters.

